# Host and pathogen drivers of infection-induced changes in social aggregation behavior

**DOI:** 10.1101/2022.05.17.492254

**Authors:** Valéria Romano, Amy Lussiana, Katy M. Monteith, Andrew J.J. MacIntosh, Pedro F. Vale

## Abstract

Identifying how infection modifies host behaviours that determine social contact networks is important for understanding heterogeneity in infectious disease dynamics. Here, we investigate whether group social behaviour is modified during bacterial infection in *Drosophila melanogaster*, an established system for behavioural genetics, according to pathogen species, infectious dose, host genetic background and sex. We find that systemic infection with four different bacterial species results in a reduction in the mean pairwise distance within infected flies, and that the extent of this change depends on the infectious dose in a pathogen-specific way. In the presence of infected conspecifics, susceptible flies also tended to aggregate throughout time, but did not show any evidence of avoiding infected flies. We also observed genetic- and sex-based variation in social aggregation, with infected female flies aggregating more closely than infected males. In general, our results confirm that bacterial infection induces changes in fruit fly behaviour across a range of pathogen species, but also highlight that these effects vary between fly genetic backgrounds and can be sex-specific. We discuss possible explanations for sex differences in social aggregation and their consequences for individual variation in pathogen transmission.

## 1. BACKGROUND

Understanding how infection modifies the group behaviour, thereby altering social connectivity and transmission dynamics, is a central focus of infectious disease research [1–5]. We can consider several types of behavioural responses to infection [6,7]. Infection avoidance is the first line of behavioural defence, where hosts modify their behaviour if they perceive an infection risk in their environment or from conspecifics [8–11]. This may include spatial or habitat avoidance [12,13], trophic avoidance [11,14,15] and social avoidance [11,16]. Nevertheless, it is rarely possible to completely avoid infection, as many common infection routes involve activities that are central to organismal physiology and fitness, including foraging and feeding. Once infected, as part of a generalized sickness response, individuals may actively self-isolate or due to their lethargic behaviour, engage in fewer social interactions [17–19], while uninfected individuals may also actively avoid those showing signals of infection[8,9,20]. Altogether, this variation in social behaviour drives the likelihood of pathogen transmission [2,21].

The extent to which hosts modify their behaviour during infection is likely to depend on their environmental and social contexts [22–24], as well as on host and pathogen genetic factors [25–27]. It is therefore important to investigate the effect of different sources of variation in infection-induced changes in insect social behaviour. The fruit fly *Drosophila melanogaster* is particularly powerful model to address this question due to its genetic tractability and its extensive use as a model of host-pathogen interactions and behavioural ecology and genetics [22,27–29]. Here, we investigate how the behavioural response to infection in Drosophila is modified by pathogen species and infectious dose, or host genetic background and sex.

In a first experiment we focus on pathogen sources of variation and ask how social aggregation behaviour changes over time when flies are exposed to either low or high doses of different bacterial pathogens. We used social groups comprised of both infected and susceptible individuals, which allowed us to test how infection affects the behaviour of infected flies, how the presence of infected flies affects the behaviour of susceptible flies, and whether there is any evidence that healthy flies show avoidance behaviour towards infected conspecifics. In a second experiment we inquire how host genetic background generates differences in social aggregation following infection, and how these effects differ between males and females.

## 2. RESULTS

### 2.1. Pathogen drivers of social aggregation

Our analysis showed a significant effect of pathogen species on the mean pairwise distance within infected flies, with a marginal trend for an interaction between dose and pathogen (Table 1, Figure 1A-B). This interaction is likely driven by flies infected with low dose (OD = 0.01) of *P. entomophila* (p = 0.0005, least square means, t ratio = -4.60) and high dose (OD = 0.1) of *E. faecalis* (p = 0.005, t ratio = -4.04) and *S. marcescens* (p = 0.04, t ratio = -3.45) when compared to control uninfected flies. We also found a marginal difference between flies infected with low dose of *S. marcescens* and controls (p = 0.08, t ratio = -3.26). When comparing the overall rate of social aggregation within infected flies to uninfected control flies, we observed that infection with almost all tested pathogens resulted in a reduction in mean pairwise distance when compared to controls: Low dose (OD = 0.01) = 1.25 mm for *E. faecalis* (Post-hoc Dunnett’s test, p ≤ 0.05), 2.61 mm for *P. entomophila* (p < 0.001), 1.05 mm for *P. rettgeri* (p = 0.11), and 1.78 mm for *S. marcescens* (p < 0.01). High dose (OD = 0.1) = 2.19 mm for *E. faecalis* (p < 0.001), 1.47 mm for *P. entomophila* (p ≤ 0.05), 1.63 mm for *P. rettgeri* (p ≤ 0.01), and 1.76 mm for *S. marcescens* (p < 0.01).

**Table 1.**
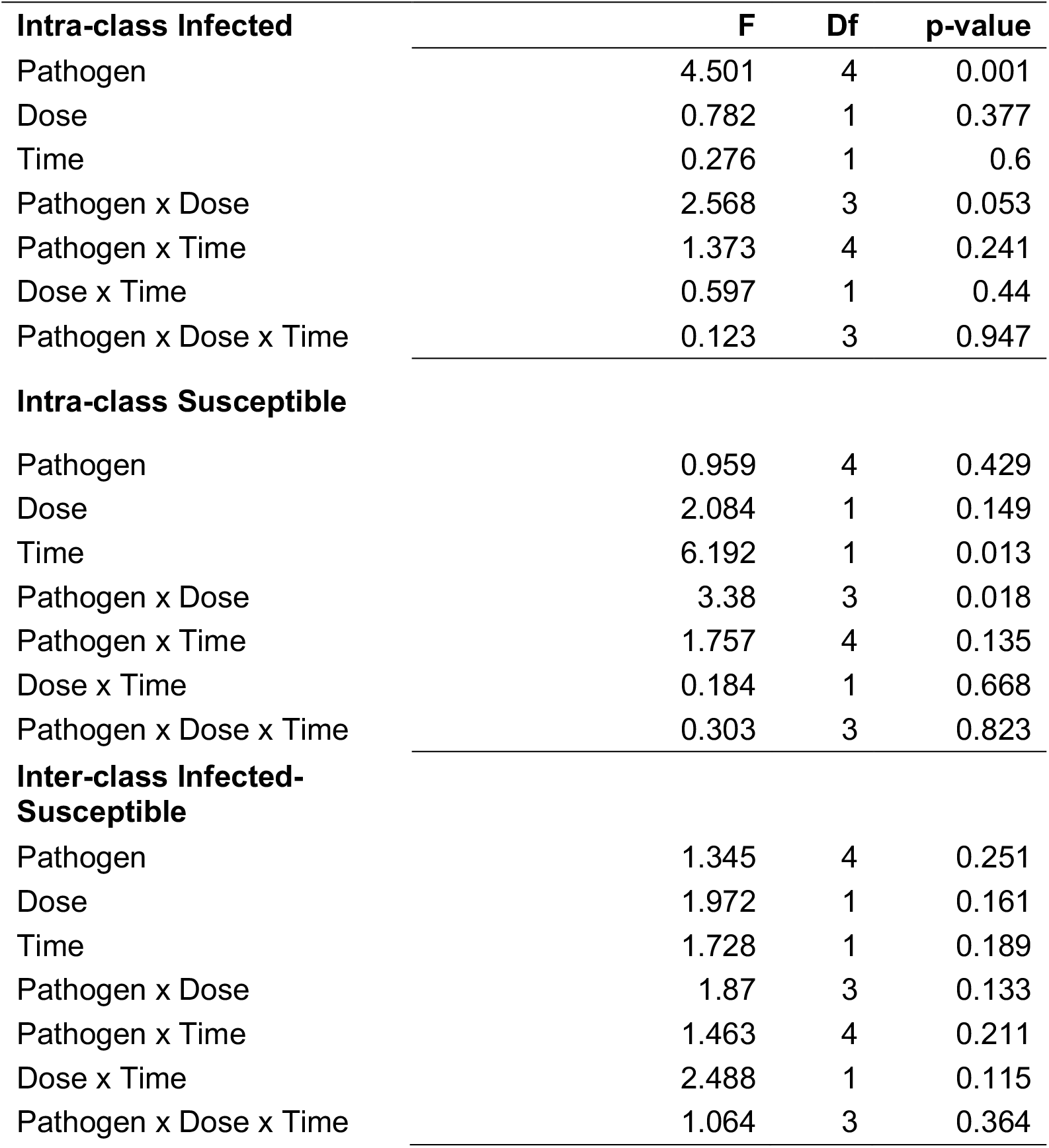
Outputs for ANOVA performed on social aggregation testing (A) intra-class pairwise distance within infected flies, (B) intra-class pairwise distance within susceptible flies, (C) inter-class pairwise distance between infected and susceptible flies.

**Figure 1.**
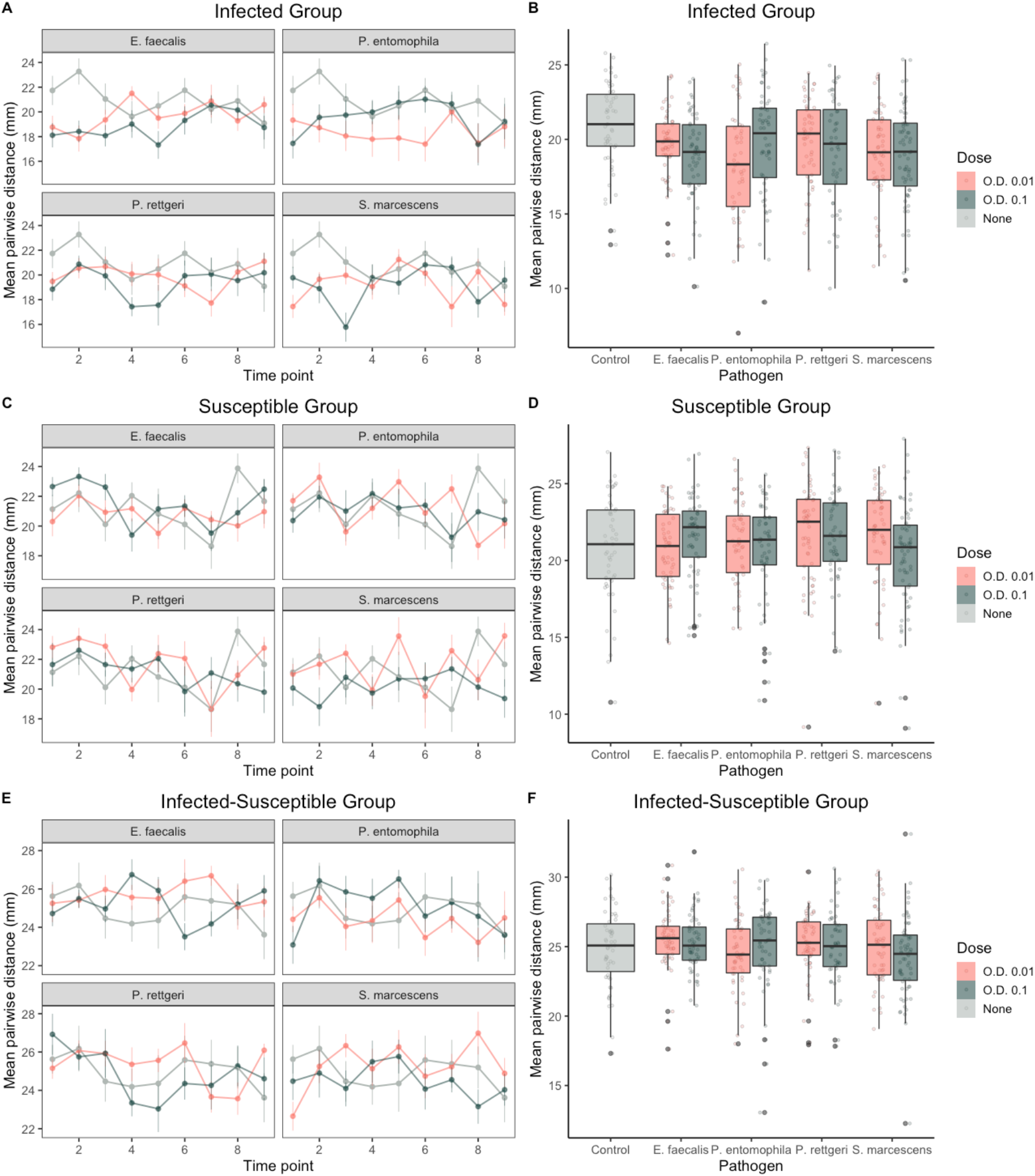
Mean pairwise distance in millimetres (mm) when considering (A, B) intra-class distance within infected flies, (C, D) intra-class distance within susceptible flies, (E, F) inter-class distance between infected and susceptible flies, of both low (O.D. 0.01) and high (O.D. 0.1) doses. A, C and E show the mean pairwise distance (mm) ± standard error (SE). B, D and F, show the mean pairwise distance (mm) for each pathogen and dose, averaged across all timepoints.

Among the subgroup of susceptible flies, we observed a reduction in the pairwise distance over the course of the experiment (Table 1, Time effect, p=0.013) and an interaction between dose and pathogen (Table 1, p=0.018, Figure 5C-D), which was marginally driven by flies infected with low and high doses of *P. rettgeri* and *S. marcescens*, respectively (p = 0.06, least square means, t ratio = 3.32). We did not observe any difference between the overall aggregation pattern of susceptible flies when compared to control groups: *E. faecalis* (Post-hoc Dunnett’s test, D1: p = 0.98; D2: p= 0.65), *P. entomophila* (D1: p = 0.96; D2: p = 1), *P. rettgeri* (D1: p = 0.32, D2: p = 0.97), and *S. marcescens* (D1: p = 0.59, D2: p = 0.42). We did not find any effect of pathogen, dose and/or time when testing the inter-class distance between infected and susceptible flies (Table 1, figure 1E-F), providing no evidence of social avoidance between susceptible and infectious flies in our experiments.

### 2.2. Host drivers of social aggregation

In a second experiment we tested whether social aggregation following systemic *P. entomophila* infection differs between flies of different genetic backgrounds and sex. We found that social aggregation is explained by host DGRP line (Figure 2A, Table 2, Line effect, p=0.001) and that patterns of social aggregation differed between males and females (Table 2, sex effect, p=0.03). We also observed a significant interaction between sex and infection status (Table 2, p=0.026, Figure 2B). While male and female flies have near identical NND aggregation in the absence of infection (p= 1, least square means, t= 0.15), infected females aggregated more closely than infected males by 1.15 mm (p = 0.01, t= - 3.04, Figure 2B). This sex difference in post-infection aggregation was observed regardless of DGRP line (there was no significant line x sex x infection interaction, Table 2).

**Figure 2.**
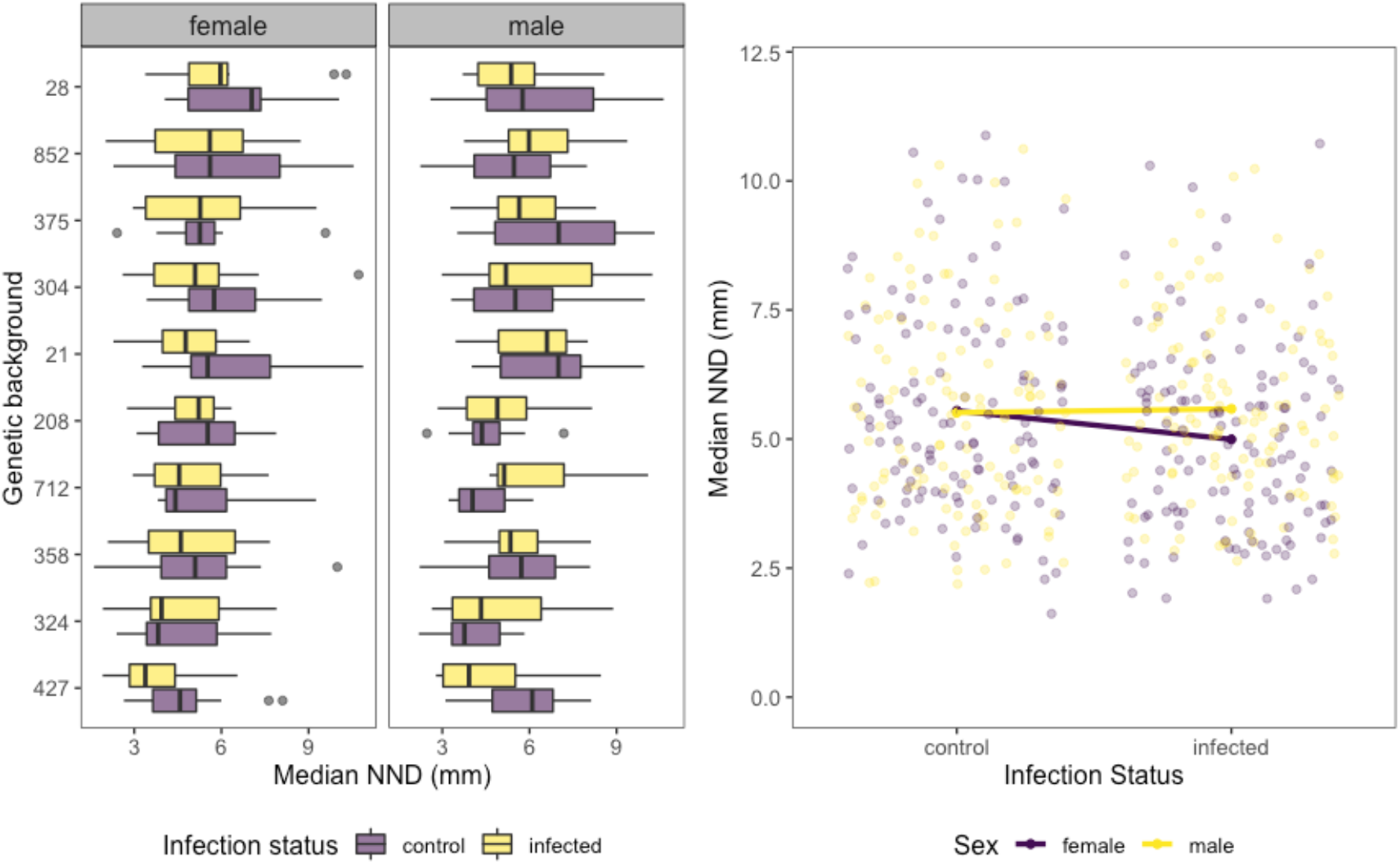
A. Boxplots showing the nearest neighbour distance (NND) in millimetres (mm) for males and females (uninfected and infected) among DGRP lines. Grey datapoints indicate outliers. B. Interaction plot of infection status and sex, based on median NND in millimetres (mm).

**Table 2.**
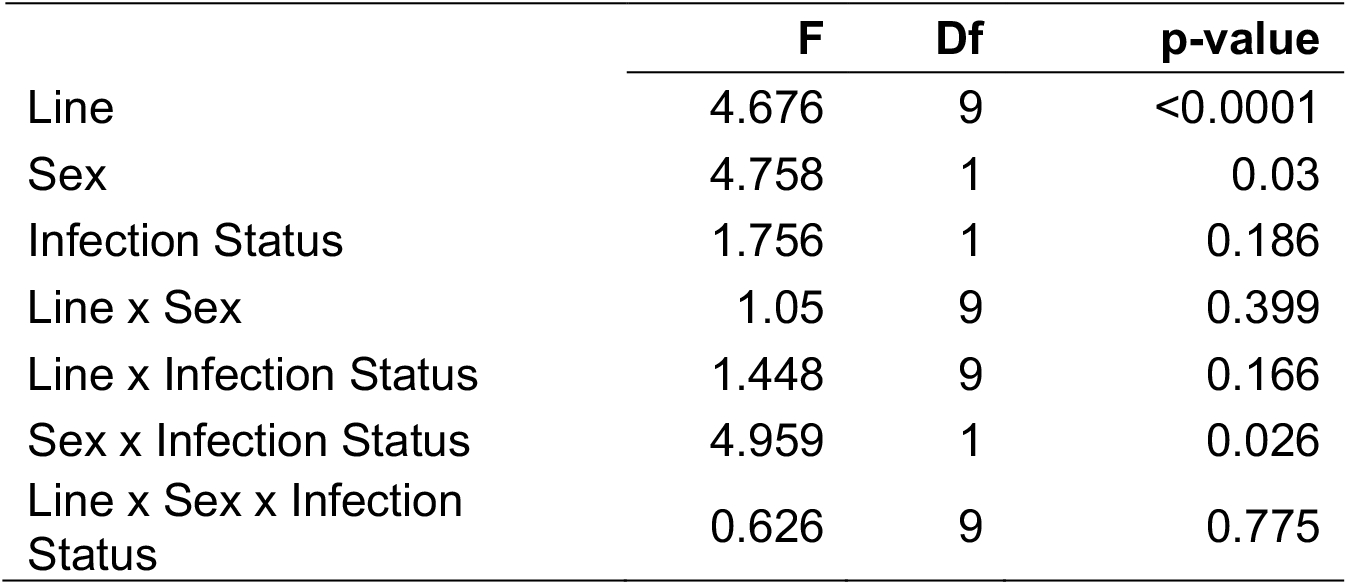
Output for ANOVA performed on social aggregation testing the influence of male and female flies of 10 DGRP lines.

|  | **F** | **Df** | **p-value** |
| --- | --- | --- | --- |
| Line | 4.676 | 9 | <0.0001 |
| Sex | 4.758 | 1 | 0.03 |
| Infection Status | 1.756 | 1 | 0.186 |
| Line x Sex | 1.05 | 9 | 0.399 |
| Line x Infection Status | 1.448 | 9 | 0.166 |
| Sex x Infection Status | 4.959 | 1 | 0.026 |
| Line x Sex x Infection Status | 0.626 | 9 | 0.775 |

## 3. Discussion

Social avoidance of infection is a widespread mechanism of defense in the animal kingdom[9,30]. Sick individuals may decrease social connectivity due to lethargic behavior or actively self-isolate [16,31], but they can also be avoided by health individuals to avoid direct routes of infection [9,20,32]. This social behavioural flexibility leads to detectable changes in the group social structure, which affects the risk of contagion among individuals [2,33]. In this study, we observed increased aggregation (shorter distances) within female infected flies – which may be due to a sickness response - but we did not find evidence that infected and susceptible flies tend to avoid each other.

Distinct ways of modifying social aggregation have been described in different social insects and may occur due to host’s social context (e.g, sex ratio,[23], alteration of feeding patterns [15], or changes in oviposition site choice [11]). An additional source of changes in infected host behavior, which we did not explore in the current study, is that pathogens can often manipulate the behavior of their hosts to increase the likelihood of transmission[7,34,35]. One relevant example relates to the increased production of attraction pheromones in flies infected with *P. entomophila*, resulting in increased aggregation between healthy and infected flies [36]. It is unclear if the increased aggregation in females we observed could have been mediated by similar pathogen-derived effects.

Regarding sex responses during infection, these appear to be pathogen specific. While this study found increased aggregation of female flies infected with pathogenic bacteria, other work identified sex differences in the opposite direction during virus infection, where males infected with Drosophila C virus aggregated further apart, with no apparent change in female social behaviour following DCV infection [25]. Sex differences in behavioural responses to infection have been described in multiple studies, and specifically in Drosophila often relate to grooming and sleeping behaviours [37,38]. A recent analysis of 59 F1-hybrids derived from the DGRP panel (the same panel of flies used here) also reported little correlation between the sociability of male and female flies [27].

One possible explanation is that sex differences are a consequence of sex-based costs of social aggregation [39]. Given that males usually display costly aggressive behaviours [40], avoiding aggregating closely when infected may also avoid the costs of aggressive encounters, while saving resources for immune deployment [41]. Female flies, however, employ generally less costly aggressive behaviours [42,43]. Differences in social aggregation costs could therefore explain why infected females aggregate more closely than males, and maintaining or augmenting sociality during infection has been suggested to reduce the impact of infection in some systems [4].

We also found that genetic background strongly influences social aggregation in fruit flies. This results confirms previous findings [25,27,44], where sociality exhibits moderate broad sense heritability (*H*^2^ = 0.21-0.24)[27], and readily responds to directional selection [45]. This large variation is to be expected for a polygenic trait such as sociality [27], and is not just characteristic of insects, as genetic background has been also found to influence social behaviours in humans and other mammalian species [9,46].

In summary, we find that flies modify their social behaviour following bacterial infection. These differences were pathogen and dose dependent, and for at least one pathogen species, this response was sexually dimorphic, with infected females aggregating more closely than infected males. Our work therefore contributes to further our understanding of this important driver of infection dynamics and of the ecology and evolution of both hosts and pathogens [2,4,32].

## 4. MATERIAL AND METHODS

### 4.1 Fly lines and bacterial strains

In experiment 1 (pathogen variation), we used female flies from a large outbred population. originally derived from DGRP (Drosophila Genetic Reference Panel). In experiment 2 (host variation), we used male and female flies from ten DGRP lines (RAL-208, RAL-852, RAL-427, RAL-304, RAL-21, RAL-375, RAL-28, RAL-324, RAL-358, RAL-712) selected to include a range of sociality scores (Anderson et al., 2015). Detailed rearing conditions are provided in Supplementary material.

### 4.2. Bacterial strains and culture

In experiment 1 we established systemic infections with one of four species of bacterial pathogen with well-described pathology in *D. melanogaster*: *Enterococcus faecalis, Pseudomonas entomophila, Serratia marcescens* DB11, and *Providencia rettgeri*. In experiment 2 we used a single bacterial fly pathogen, *P. entomophila* at OD = 0.1. Detailed culture conditions are provided in Supplementary material.

### 4.3. Experiment 1 (Pathogen variation)

Social interaction chambers consisted of 50 mm Petri dishes containing 8% sugar-agar medium. In total we set up 24-replicate social groups for each pathogen and dose (N= 192), plus 24 control groups. Flies were anaesthetised using light CO_2_ and infected in the mesopleuron with one of four bacterial pathogens at OD 0.1 or 0.01 using a 0.14 mm diameter stainless steel pin. The experiment was blocked over 4 consecutive days, with chambers including all different treatments spread across each block day. Infections were always carried out between 10am-2pm on each day. Each Petri dish contained six uninfected, susceptible female flies and six female flies infected with a specific bacterial pathogen at a specific dose. Infected flies were marked with red fluorescent powder on the prothorax and the underside of the abdomen using a cotton bud. Control plates were also setup containing twelve uninfected individuals, with half marked as above. Flies were allowed an hour of recovery from the systemic infection and marking before being re-anaesthetised using light CO_2_ and added to the social interaction chambers. Thirty minutes were allowed for habituation before photos of the groups were taken every 30 minutes until 4 hours post-infection. Pictures in which the infected and susceptible individuals could be reliably identified were processed in ImageJ, where we estimated coordinates of each individual. Social aggregation was then measured in R (R Core Team, 2020) using the *pairdist* function in the *spatstat* package. The pairwise distance between each pair of flies within a dish was used to calculate three sociality metrics per dish: (i) the mean pairwise distance between infected flies, which is relevant to evaluate changes in aggregation due to sickness behaviour; (ii) the mean pairwise distance between susceptible flies, and (iii) the mean pairwise distance between infected and susceptible flies, which allows to test if susceptible flies tend to avoid infected flies, when compared to the control group. Therefore, each dish resulted in two intra-class measures (within infected and within susceptible) and one inter-class measures (between infected and susceptible).

### 4.4. Experiment 2 (Host variation)

For each of the ten fly lines, we set up single-sex groups of flies, divided into infected and control, with each fly line-sex-treatment was replicated 11-12 times, for a total of 466 social aggregation assays. Each group consisted of 12 flies systemically infected with *P. entomophila* (or sterile LB medium for uninfected control groups) using a stainless pin. Following infection, flies were lightly anaesthetised with CO_2_ and transferred to 55 mm Petri dishes containing agar. After a 30-minute habituation period, photographs were taken. Here we used the median nearest neighbour distance (NND) of each group as a measure of social aggregation [25,44]. Individual fly positions in each image were marked in the middle of the fly thorax using Fiji (**F**iji **I**s **J**ust **I**mageJ), and the nearest neighbour distances between each pair of flies was calculated using the ‘NND’ plugin within the software Fiji (Schindelin et al, 2012).

### 4.5. Statistical analysis

All raw data and analysis R code are available at https://doi.org/10.5281/zenodo.6554320. Data from experiment 1 was analysed using linear mixed effects models, separately for each social class (i.e., within infected, within susceptible, between infected and susceptible). We used the mean pairwise distance as the response factor, pathogen, dose and time as predictor variables, and day of assay as a random effect. For experiment 2, we used a linear mixed effects model with the log_10_ of median NND as the response variable, line, sex and infection status as predictors, and day of the assay as a random effect. All possible interactions between line, sex and infection status were included. A more detailed description of the analysis can be found in Supplementary material.

## 5. Acknowledgements

We are grateful to A. Reid for media preparation and J. Boyle for help in collecting data for experiment 2. We thank members of the Regan and Obbard Labs for helpful comments. This work was partially funded by a Society in Science Branco Weiss Fellowship, and a Chancellor’s Fellowship, both awarded to P Vale.

## Supplementary material for

**Figure S1.**
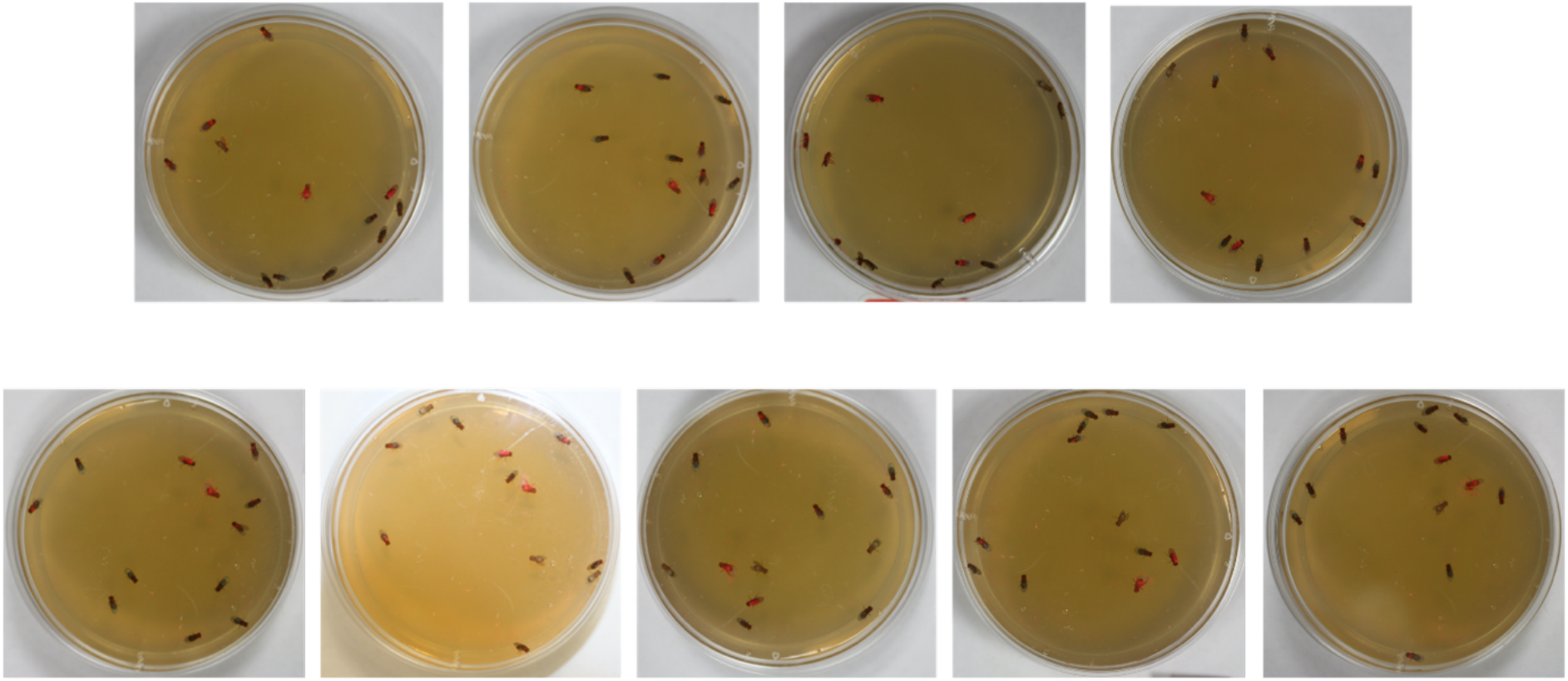
Examples of pictures taken of marked flies every 30 mins.

**Figure S2.**
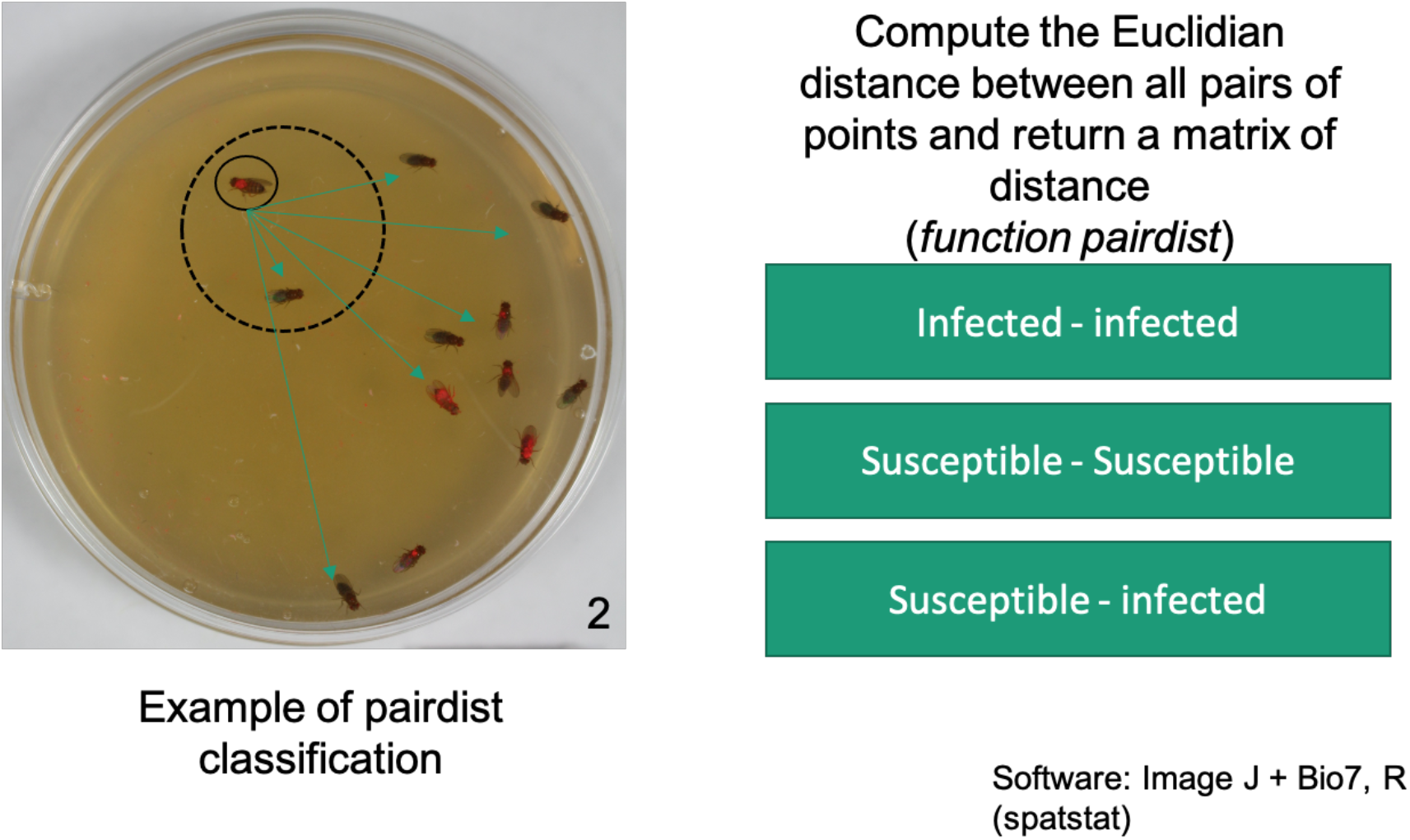
Examples of pairwise distance calculation.

## Supplementary methods

### Flies and rearing conditions

In experiment 1 (pathogen variation), we used age-matched and mated female flies from a large outbred population. This outbred population was originally derived from DGRP (Drosophila Genetic Reference Panel) population (Mackay et al., 2012) and was established by randomised pairwise crosses between virgin males and females from 113 DGRP lines to maximise genetic diversity. All lines were cleared of Wolbachia before outcrossing. Prior to experimentation the outcrossed population had been maintained for at least 43 generations at 25 ± 1 °C with a 12 L : 12 D cycle in bottles containing standard cornmeal-sugar medium. Before the experiment females were left in stock bottles for three days post-eclosion to allow mating, sorted into cohorts of 80 individuals and maintained under controlled light and temperature as above until required for experimentation. In experiment 2 (host variation), flies were maintained as above, but in vials. We used age-matched 2-4 day-old, mated male and female flies from ten DGRP lines (RAL-208, RAL-852, RAL-427, RAL-304, RAL-21, RAL-375, RAL-28, RAL-324, RAL-358, RAL-712) selected to include a range of sociality scores (Anderson et al., 2015).

### Bacterial strains and culture

In experiment 1 (pathogen variation) we established systemic infections with one of four species of bacterial pathogen with well-described pathology in *D. melanogaster*: *Enterococcus faecalis, Pseudomonas entomophila, Serratia marcescens* DB11, and *Providencia rettgeri* (Chandler et al., 2011; Vodovar et al., 2005; Corby-Harris et al., 2007; Cox & Gilmore, 2007). Frozen stocks were isolated from a single bacterial colony, aliquoted and stored in a 25% glycerol suspension at -70°C. Fresh bacterial cultures were generated daily by overnight culturing for ∼14 hrs in 10 mL Luria-Bertani (LB) broth growth medium at 30 °C, 140 rpm and aerobic conditions. When the bacterial culture had reached the mid-log phase of growth (optical density: OD ∼ 0.8) the bacterial suspension was serially diluted in LB broth to OD = 0.1 (high dose) and OD = 0.01 (low dose). In experiment 2 (host variation) we wanted to focus on host variation, and so to minimise other sources of variation we decided used a single bacterial fly pathogen, *P. entomophila*, prepared as described above, and subsequently diluted with LB medium to OD = 0.1.

### Statistical analysis

All raw data and analysis code are available at https://doi.org/10.5281/zenodo.6554320. All statistical analyses and graphics were carried out and produced in R 4.0.5 using the ggplot2 [47], car [48], dplyr [49], lme4[50], lmerTest [51], emmeans [52] and DescTools (x) packages. Data from experiment 1 was analysed using linear mixed effects models, one per each class distance (i.e., within infected, within susceptible, between infected and susceptible). We used the mean pairwise distance as the response factor, pathogen, dose and time as predictor variables, and day of assay as a random effect. All possible interactions between pathogen, dose and time were included. For experiment 2, median NND was log-transformed to achieve a normal distribution of residuals. We then used a linear mixed effects model with the log_10_ of median NND as the response variable, line, sex and infection status as predictors, and day of the assay as a random effect. All possible interactions between line, sex and infection status were included. We used ANOVA and adjusted least-square means to test the significance of fixed effects and perform post hoc comparisons among groups when interactions were significant. We also applied Dunnett tests to compare the overall rate of social aggregation between the control and infected flies. Model assumptions (homogeneity of variance, normality of residuals and random effects, collinearity for additive models,) were tested and found to be fulfilled.

## References

1. Barron D, Gervasi S, Pruitt J, Martin L. 2015 Behavioral competence: how host behaviors can interact to influence parasite transmission risk. Curr. Opin. Behav. Sci. 6, 35–40. (doi:10.1016/j.cobeha.2015.08.002)

2. Romano V, MacIntosh AJJ, Sueur C. 2020 Stemming the Flow: Information, Infection, and Social Evolution. Trends Ecol. Evol. 35, 849–853. (doi:10.1016/j.tree.2020.07.004)

3. White LA, Siva-Jothy JA, Craft ME, Vale PF. 2020 Genotype and sex-based host variation in behaviour and susceptibility drives population disease dynamics. Proc. R. Soc. B Biol. Sci. 287, 20201653. (doi:10.1098/rspb.2020.1653)

4. Hawley DM, Gibson AK, Townsend AK, Craft ME, Stephenson JF. 2021 Bidirectional interactions between host social behaviour and parasites arise through ecological and evolutionary processes. Parasitology 148, 274–288. (doi:10.1017/S0031182020002048)

5. Craft ME. 2015 Infectious disease transmission and contact networks in wildlife and livestock. Philos. Trans. R. Soc. B Biol. Sci. 370, 20140107. (doi:10.1098/rstb.2014.0107)

6. de Roode JC, Lefèvre T. 2012 Behavioral Immunity in Insects. Insects 3, 789–820. (doi:10.3390/insects3030789)

7. Vale PF, Siva-Jothy J, Morrill A, Forbes M. 2018 The influence of parasites on insect behavior. In Insect Behavior: from mechanisms to ecological and evolutionary consequences, pp. 274–291. OUP Oxford.

8. Behringer DC, Butler MJ, Shields JD. 2006 Ecology: Avoidance of disease by social lobsters. Nature 441, 421–421. (doi:10.1038/441421a)

9. Curtis VA. 2014 Infection-avoidance behaviour in humans and other animals. Trends Immunol. 35, 457–464. (doi:10.1016/j.it.2014.08.006)

10. Eakin L, Wang M, Dwyer G. 2015 The effects of the avoidance of infectious hosts on infection risk in an insect-pathogen interaction. Am. Nat. 185, 100–112. (doi:10.1086/678989)

11. Siva-Jothy JA, Monteith KM, Vale PF. 2018 Navigating infection risk during oviposition and cannibalistic foraging in a holometabolous insect. Behav. Ecol. Off. J. Int. Soc. Behav. Ecol. 29, 1426–1435. (doi:10.1093/beheco/ary106)

12. Amano H, Hayashi K, Kasuya E. 2008 Avoidance of egg parasitism through submerged oviposition by tandem pairs in the water strider, Aquarius paludum insularis (Heteroptera: Gerridae). Ecol. Entomol. 33, 560–563. (doi:10.1111/j.1365-2311.2008.00988.x)

13. Mierzejewski MK, Horn CJ, Luong LT. 2019 Ecology of fear: environment-dependent parasite avoidance among ovipositing Drosophila. Parasitology 146, 1564–1570. (doi:10.1017/S0031182019000854)

14. Parker BJ, Elderd BD, Dwyer G. 2010 Host behaviour and exposure risk in an insect– pathogen interaction. J. Anim. Ecol. 79, 863–870. (doi:10.1111/j.1365-2656.2010.01690.x)

15. Kobler JM, Rodriguez Jimenez FJ, Petcu I, Grunwald Kadow IC. 2020 Immune Receptor Signaling and the Mushroom Body Mediate Post-ingestion Pathogen Avoidance. Curr. Biol. CB 30, 4693-4709.e3. (doi:10.1016/j.cub.2020.09.022)

16. Bos N, Lefèvre T, Jensen AB, D’ettorre P. 2012 Sick ants become unsociable. J. Evol. Biol. 25, 342–351. (doi:10.1111/j.1420-9101.2011.02425.x)

17. Hart BL. 1988 Biological basis of the behavior of sick animals. Neurosci. Biobehav. Rev. 12, 123–137. (doi:10.1016/s0149-7634(88)80004-6)

18. Lopes PC, French SS, Woodhams DC, Binning SA. 2021 Sickness behaviors across vertebrate taxa: proximate and ultimate mechanisms. J. Exp. Biol. 224, jeb225847. (doi:10.1242/jeb.225847)

19. Kazlauskas N, Klappenbach M, Depino AM, Locatelli FF. 2016 Sickness Behavior in Honey Bees. Front. Physiol. 7. (doi:10.3389/fphys.2016.00261)

20. Meisel JD, Kim DH. 2014 Behavioral avoidance of pathogenic bacteria by Caenorhabditis elegans. Trends Immunol. 35, 465–470. (doi:10.1016/j.it.2014.08.008)

21. Stockmaier S, Stroeymeyt N, Shattuck EC, Hawley DM, Meyers LA, Bolnick DI. 2021 Infectious diseases and social distancing in nature. Science 371, eabc8881. (doi:10.1126/science.abc8881)

22. Dukas R. 2020 Natural history of social and sexual behavior in fruit flies. Sci. Rep. 10, 21932. (doi:10.1038/s41598-020-79075-7)

23. Keiser CN, Rudolf VHW, Sartain E, Every ER, Saltz JB. 2018 Social context alters host behavior and infection risk. Behav. Ecol. 29, 869–875. (doi:10.1093/beheco/ary060)

24. Lopes PC, Adelman J, Wingfield JC, Bentley GE, Wingfield JC, Lopes PC, Adelman J. 2012 Social context modulates sickness behavior. Behav. Ecol. Sociobiol. 66, 1421– 1428. (doi:10.1007/s00265-012-1397-1)

25. Siva-Jothy JA, Vale PF. 2019 Viral infection causes sex-specific changes in fruit fly social aggregation behaviour. Biol. Lett. 15, 20190344. (doi:10.1098/rsbl.2019.0344)

26. Keiser CN, Rudolf VHW, Luksik MC, Saltz JB. 2020 Sex differences in disease avoidance behavior vary across modes of pathogen exposure. Ethology 126, 304– 312. (doi:10.1111/eth.12969)

27. Scott AM, Dworkin I, Dukas R. 2018 Sociability in Fruit Flies: Genetic Variation, Heritability and Plasticity. Behav. Genet. 48, 247–258. (doi:10.1007/s10519-018-9901-7)

28. Dubnau J. 2014 Behavioral Genetics of the Fly (Drosophila Melanogaster). Cambridge University Press.

29. Buchon N, Silverman N, Cherry S. 2014 Immunity in Drosophila melanogaster — from microbial recognition to whole-organism physiology. Nat. Rev. Immunol. 14, 796–810. (doi:10.1038/nri3763)

30. Curtis V, de Barra M, Aunger R. 2011 Disgust as an adaptive system for disease avoidance behaviour. Philos. Trans. R. Soc. Lond. B. Biol. Sci. 366, 389–401. (doi:10.1098/rstb.2010.0117)

31. Cremer S, Armitage SAO, Schmid-Hempel P. 2007 Social Immunity. Curr. Biol. 17, R693–R702. (doi:10.1016/j.cub.2007.06.008)

32. Buck JC, Weinstein SB, Young HS. 2018 Ecological and Evolutionary Consequences of Parasite Avoidance. Trends Ecol. Evol. 33, 619–632. (doi:10.1016/j.tree.2018.05.001)

33. Lopes PC, Block P, König B. 2016 Infection-induced behavioural changes reduce connectivity and the potential for disease spread in wild mice contact networks. Sci. Rep. 6, 31790. (doi:10.1038/srep31790)

34. Poulin R, Brodeur J, Moore J. 1994 Parasite Manipulation of Host Behaviour: Should Hosts Always Lose? Oikos 70, 479–484. (doi:10.2307/3545788)

35. van Houte S, Ros VID, van Oers MM. 2013 Walking with insects: molecular mechanisms behind parasitic manipulation of host behaviour. Mol. Ecol. 22, 3458– 3475. (doi:10.1111/mec.12307)

36. Keesey IW, Koerte S, Khallaf MA, Retzke T, Guillou A, Grosse-Wilde E, Buchon N, Knaden M, Hansson BS. 2017 Pathogenic bacteria enhance dispersal through alteration of Drosophila social communication. Nat. Commun. 8, 265. (doi:10.1038/s41467-017-00334-9)

37. Yanagawa A, Guigue AMA, Marion-Poll F. 2014 Hygienic grooming is induced by contact chemicals in Drosophila melanogaster. Front. Behav. Neurosci. 8.

38. Vale PF, Jardine MD. 2015 Sex-specific behavioural symptoms of viral gut infection and Wolbachia in Drosophila melanogaster. J. Insect Physiol. 82, 28–32. (doi:10.1016/j.jinsphys.2015.08.005)

39. Kelly CD, Stoehr AM, Nunn C, Smyth KN, Prokop ZM. 2018 Sexual dimorphism in immunity across animals: a meta-analysis. Ecol. Lett. 21, 1885–1894. (doi:10.1111/ele.13164)

40. Chen S, Lee AY, Bowens NM, Huber R, Kravitz EA. 2002 Fighting fruit flies: a model system for the study of aggression. Proc. Natl. Acad. Sci. U. S. A. 99, 5664–5668. (doi:10.1073/pnas.082102599)

41. Adamo SA, Gomez-Juliano A, LeDue EE, Little SN, Sullivan K. 2015 Effect of immune challenge on aggressive behaviour: how to fight two battles at once. Anim. Behav. 105, 153–161. (doi:10.1016/j.anbehav.2015.04.018)

42. Ueda A, Kidokoro Y. 2002 Aggressive behaviours of female Drosophila melanogaster are influenced by their social experience and food resources. Physiol. Entomol. 27, 21–28. (doi:10.1046/j.1365-3032.2002.00262.x)

43. Nilsen SP, Chan Y-B, Huber R, Kravitz EA. 2004 Gender-selective patterns of aggressive behavior in Drosophila melanogaster. Proc. Natl. Acad. Sci. 101, 12342– 12347. (doi:10.1073/pnas.0404693101)

44. Anderson BB, Scott A, Dukas R. 2016 Social behavior and activity are decoupled in larval and adult fruit flies. Behav. Ecol. 27, 820–828. (doi:10.1093/beheco/arv225)

45. Scott AM, Dworkin I, Dukas R. 2022 Evolution of sociability by artificial selection*. Evolution 76, 541–553. (doi:10.1111/evo.14370)

46. Anacker AMJ, Beery AK. 2013 Life in groups: the roles of oxytocin in mammalian sociality. Front. Behav. Neurosci. 7, 185. (doi:10.3389/fnbeh.2013.00185)

47. Wickham H. 2009 ggplot2: Elegant Graphics for Data Analysis. Ggplot2 Elegant Graph. Data Anal., 1–212. (doi:10.1007/978-0-387-98141-3)

48. Fox J;, Weisberg S. 2011 An R Companion to Applied Regression, Second Edition.

49. Wickham H, Francois R, Henry L, Müller K. 2019 dplyr: A Grammar of Data Manipulation. Ggplot2 Elegant Graph. Data Anal., 1–212. (doi:10.1007/978-0-387-98141-3)

50. Bates D et al. 2018 lme4: Linear Mixed-Effects Models using ‘Eigen’ and S4. See https://CRAN.R-project.org/package=lme4.

51. Kuznetsova A, Brockhoff PB, Christensen RHB. 2017 lmerTest Package: Tests in Linear Mixed Effects Models. J. Stat. Softw. 82. (doi:10.18637/jss.v082.i13)

52. Lenth RV, Buerkner P, Herve M, Love J, Miguez F, Riebl H, Singmann H. 2022 emmeans: Estimated Marginal Means, aka Least-Squares Means. See https://CRAN.R-project.org/package=emmeans.

